# Mitochondrial ClpX activates an essential metabolic enzyme through partial unfolding

**DOI:** 10.1101/327056

**Authors:** Julia R. Kardon, Jamie A. Moroco, John R. Engen, Tania A. Baker

**Affiliations:** Department of Biology, Massachusetts Institute of Technology, Cambridge, MA 02139, USA; Howard Hughes Medical Institute, Massachusetts Institute of Technology, Cambridge, MA 02139, USA; Department of Chemistry and Chemical Biology, Northeastern University, Boston, MA 02115, USA

## Abstract

Mitochondria can control the activity, quality, and lifetime of their proteins with their autonomous system of chaperones, but the signals that direct substrate-chaperone interaction and outcome are poorly understood. We previously discovered that the mitochondrial AAA+ protein unfoldase ClpX (mtClpX) activates the initiating enzyme for heme biosynthesis, 5-aminolevulinic acid synthase (ALAS), by promoting incorporation of cofactor. Here, we ask how unfolding by mtClpX directs activation. We identified sequence and structural features in ALAS that position mtClpX and provide a grip for acting on ALAS. Observation of ALAS undergoing remodeling by mtClpX revealed that unfolding was limited to a subdomain extending from the mtClpX-binding site to the active site. Unfolding along this path was required for mtClpX to gate cofactor access to the ALAS active site. This targeted unfolding contrasts with the global unfolding canonically executed by ClpX homologs and suggests how substrate-chaperone interactions can direct the outcome of remodeling.

## INTRODUCTION

AAA+ protein unfoldases use the energy of ATP hydrolysis to pull apart the structures of their substrate proteins. In general, this is accomplished by gripping the substrate polypeptide with aromatic side chains of loops that protrude into the central pore of the unfoldase hexamer; ATP hydrolysis drives conformational changes in the hexamer that push these loops downward, pulling and thus mechanically unfolding the substrate from the site of engagement^1^.

Although some AAA+ unfoldases are specialized for non-proteolytic protein unfolding, this activity has been best characterized as part of protein degradation, in which the polypeptide is directly translocated into the proteolytic chamber of a co-complexed peptidase. This basic architecture and mechanism is used by eukaryotic and archaeal proteasomes and by several unfoldase-protease complexes shared among bacteria, mitochondria, peroxisomes, and chloroplasts^1–3^. One such bacterial unfoldase, ClpX, with its partner peptidase ClpP, is particularly specialized for regulatory degradation of substrates rather than protein quality control, conditionally selecting a varied repertoire of substrates to execute stress responses and cell fate decisions^4^. Accumulating evidence indicates that mitochondrial ClpX (mtClpX) acts in a similarly regulatory capacity, although only a few substrates of mtClpX have been verified and the specific regulatory consequences of its actions on mitochondrial physiology are for the most part undetermined^5–10^. How mtClpX recognizes its substrates has also not been characterized, although its sequence preferences for substrate recognition clearly diverges from that of bacterial ClpX^11^.

We previously discovered that mtClpX) promotes heme biosynthesis by acting on the first enyzme in heme biosynthesis, 5-aminolevulinate synthase (ALAS)^5^. In contrast to unfolding of a protein substrate for degradation, as ClpX homologs are best understood to do, mtClpX activates ALAS by accelerating incorporation of its cofactor, pyridoxal 5’-phosphate (PLP). This activation is non-proteolytic and requires ATP hydrolysis and intact pore loops, implicating mtClpX unfoldase activity in this unconventional activity. Here, we address how mtClpX is specifically deployed to direct activation of an enzyme. Using a peptide array of the ALAS sequence combined with protein mutagenesis and engineering, we defined a coherent mtClpX-binding site that spans the dimer interface of ALAS. This site contains separable elements that recruit mtClpX and engage mtClpX for initiation of unfolding. Using hydrogen-deuterium exchange coupled with mass spectrometry (HX MS), we observed that mtClpX induces exposure of a limited region of ALAS that extends from the mtClpX-binding site to the active site, serving as a gate to the active site. Engineered crosslinks that obstruct this path demonstrated that this remodeling is necessary for ALAS activation. Our observations describe the mechanism by which a conserved mitochondrial unfoldase activates an essential biosynthetic enzyme and provide a model for how the interactions between protein unfoldases and their substrates can be directed to widely divergent outcomes, from restricted unfolding and activation to complete unfolding and degradation.

## RESULTS

### An N-terminal sequence directs mtClpX to activate ALAS

To determine how mtClpX induces ALAS to bind PLP, we first sought to identify how mtClpX recognizes ALAS and engages it for unfolding. As an unbiased strategy, we assayed binding of mtClpX to an array of peptides representing the linear sequence of ALAS. mtClpX bound six discrete peptide sequences within ALAS (Fig. 1A, blue; Fig. S1C). Five of these sequences map to a structurally contiguous site in ALAS, consisting of an α-helix (α1) that is the most N-terminal structured element of ALAS, and a small region that is in direct hydrogen-bonding contact with α1 across the ALAS dimer interface (Fig. 1B). The sixth, most C-terminal peptide maps to a separate site about 40 Å away.

**Figure 1.**
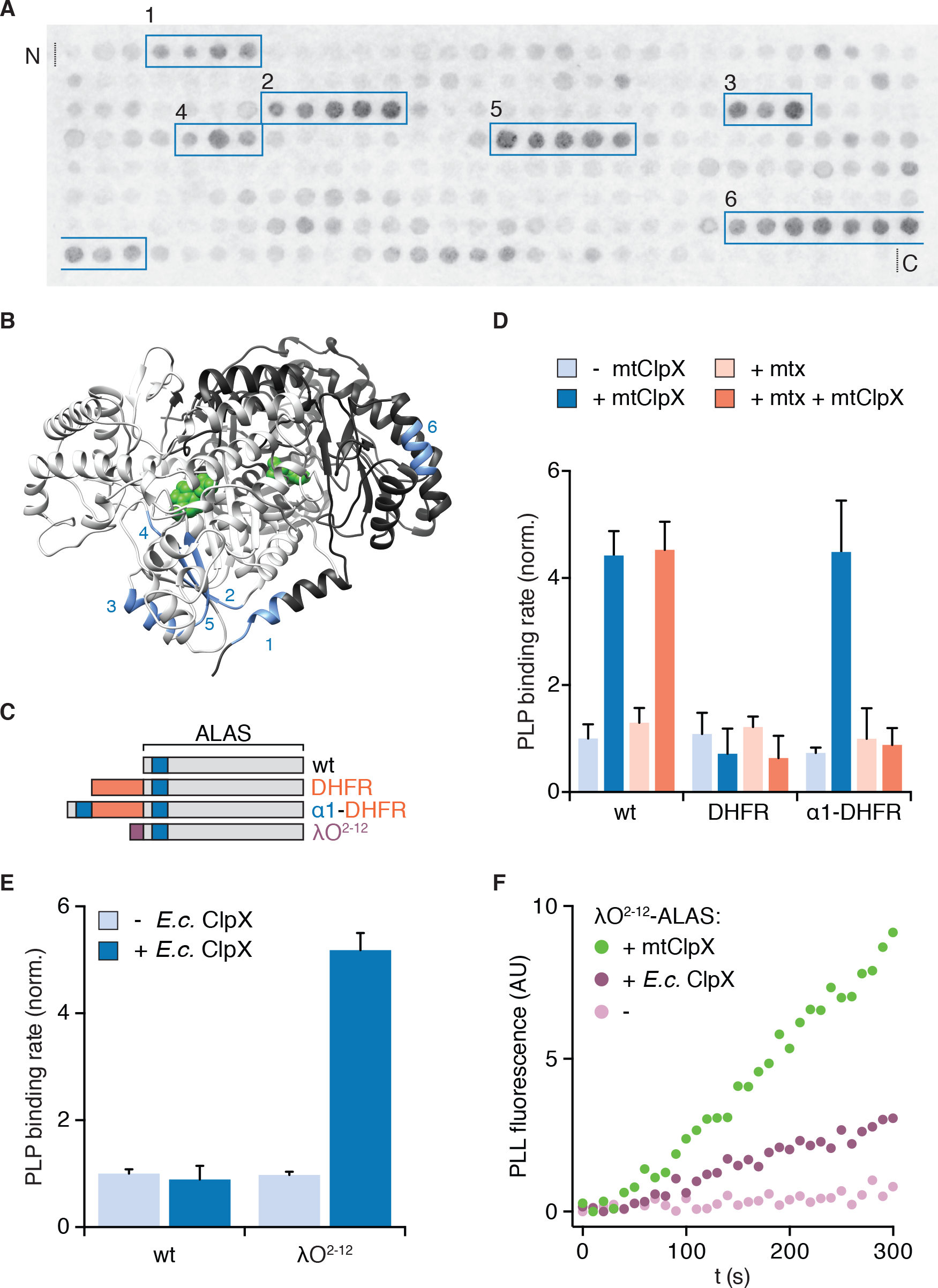
Interaction with the N-terminus of ALAS is necessary and sufficient for activation of ALAS by an unfoldase. (**A**) Peptide array of the ALAS sequence (58-548, sliding window of fifteen amino acids, shifted two amino acids towards the C-terminus with each spot, N-to C-terminus arrayed left-to-right, top-to-bottom) probed with mtClpX^E206Q^-3xFLAG and detected by far-western blot. mtClpX-binding sequences are boxed in blue. See Fig. S1C for control blot. **(B)** mtClpX-binding sequences identified by peptide blotting are mapped on one face of the structure of *S. cerevisiae* ALAS (PDB: 5TXR) in blue and numbered as in (A). Sequences were defined as the range between the two amino acids added at the beginning of the boxed region in (A) and the two amino acids removed after its end. PLP is depicted in green and the two protomers of ALAS are colored in light and dark gray, respectively. **(C)** Diagram of ALAS N-terminal variants. Blue indicates the N-terminal ClpX-binding peptide in α1, orange indicates *M. musculus* dihydrofolate reductase (DHFR), and purple indicates a degradation tag recognized by *E. coli* ClpX (residues 2-12 of the phage λO replication protein). **(D)** Rate of PLP binding to ALAS and N-terminal DHFR variants (5 μM monomer), ± mtClpX (2 μM hexamer), ± methotrexate (mtx) (30 μM). Reactions additionally contained 2 mM ATP and a regeneration system and 50 mM PLP (see Materials and Methods). PLP binding was monitored by fluorescence specific to protein-liganded PLP (ex. 434 nm, em. 515 nm). Rates were extracted by linear fits to values in the early linear phase and normalized to the rate for wildtype ALAS without methotrexate or mtClpX. p < 0.001 for suppression of mtClpX activity by DHFR fusion (DHFR-ALAS) and suppression of mtClpX activity on α1-DHFR-ALAS by methotrexate additon. (**E)** PLP binding to ALAS and λO ^2-12^-ALAS (5 μM monomer), ± *E. coli* ClpX (2 μM hexamer), assayed as in (C). p < 10^−4^ for stimulation of PLP binding to λO^2-12^-ALAS by *E. coli* ClpX. **(F)** PLP binding fluorescence traces for λO ^2-12^-ALAS, ± *E. coli* ClpX or mtClpX. Error bars represent standard deviation; n ≥ 3. P values were calculated using Student’s t-test.

Because ClpX and other AAA+ unfoldases often initiate unfolding at or near protein termini, α1 of ALAS was an appealing candidate for an initial binding site for mtClpX. α1 is also immediately adjacent in structure and in sequence to a small β-sheet (β1-3) that abuts the active site. Unfolding initiated at α1 could restructure this region to allow PLP access to the active site. Indeed, we observed that β1-3 is conformationally responsive to the presence of PLP in the active site in our crystal structure of ALAS^12^: One protomer of the ALAS dimer lost PLP during crystallization in PLP-free solvent and β1-3 of the unoccupied protomer had weaker electron density and could not be modeled.

We previously also crystallized ALAS that had been incubated with hydroxylamine, which cleaves the covalent PLP-lysine bond in the active site and converts PLP to non-covalently bound and inactive PLP-oxime (Fig. S1A)^12^. Our crystal structure from this inactive, PLP-depleted preparation contains a PLP-derived species in the active site despite gel filtration to remove free cofactor. Because our previous assays for activation of ALAS by mtClpX were performed with this hydroxylamine-treated preparation of ALAS, we sought to determine whether mtClpX exchanges inactive cofactor, facilitates cofactor binding to unoccupied active sites, or both. We determined that hydroxylamine-treated and gel-filtered ALAS remained slightly more than half-occupied with a species consistent with PLP-oxime (Fig. S1A, B). The activation of hydroxylamine-treated ALAS by mtClpX therefore represents a combination of PLP binding to unoccupied active sites and exchange of PLP-oxime for PLP. Both events are likely useful in supporting ALAS function in vivo, assisting de novo PLP binding and regenerating ALAS enzyme inhibited by damaged cofactor (for example, PMP resulting from decarboxylation-coupled transamination of PLP, or other natural errors in catalysis^13^). Both events could be similarly facilitated by unfolding by mtClpX to expose the ALAS active site.

To test the importance of the putative N-terminal mtClpX binding site of ALAS, we engineered N-terminal fusions of ALAS with dihydrofolate reductase (DHFR) (Fig. 1C). N-terminal fusion of DHFR with ALAS (DHFR-ALAS) blocked stimulation of PLP binding by mtClpX, but appending the N-terminal sequence of ALAS through α1 to DHFR-ALAS (α1-DHFR-ALAS) restored the ability of mtClpX to stimulate PLP binding. Thus, mtClpX recognizes the N-terminus of ALAS to initiate activation. Upon binding its inhibitor methotrexate, DHFR becomes extremely mechanically stable and resistant to unfolding, thus providing a conditional block to mtClpX. Addition of methotrexate blocked mtClpX activation of α1-DHFR-ALAS (Fig. 1D) indicating that unfolding from the N-terminus is required for ALAS activation. Therefore, mtClpX requires the α1 site to recognize ALAS and unfolds from the N-terminus to induce activation.

### Unfolding from the N-terminus is sufficient to activate ALAS

To test if unfolding from the N-terminus of ALAS is sufficient for activation, we attempted to direct *E. coli* ClpX to stimulate PLP binding. *E. coli* ClpX does not act on wildtype ALAS (Fig. 1E). To direct it to ALAS, we appended a recognition sequence for *E. coli* ClpX (λO^2-12^, from the *E. coli* ClpX substrate lambda phage O protein^14^) to the N-terminus of ALAS (λO^2-12^-ALAS) (Fig. 1C). *E. coli* ClpX accelerated PLP binding to λO^2-12^-ALAS at a similar rate to mtClpX with wildtype ALAS (Fig. 1E, F). Therefore, an unfoldase acting from this N-terminal site is sufficient to stimulate ALAS activation. We also tested if the short, unstructured λO^2-12^ tag might block mtClpX action at the N-terminus of ALAS as the folded DHFR domain did. We unexpectedly observed the opposite effect: λO^2-12^-ALAS was processed more rapidly than ALAS by mtClpX (Fig. 1F). This increased efficiency was not based on specific interaction of mtClpX with the λO^2-12^ peptide (Fig. S1D), suggesting that mtClpX instead responded to the unstructured extension contributed by the λO^2-12^ tag.

### mtClpX relies on an unstructured N-terminal extension for rapid activation of ALAS

The observation that an unstructured extension to the N-terminus of ALAS stimulated its activation by mtClpX led us to reexamine the native N-terminus of ALAS in the mitochondrion. Many mitochondrial proteins, including ALAS, are translated with an N-terminal targeting sequence that is cleaved after import into the mitochondrial matrix. The N-terminus of the ALAS constructs we used in our previous experiments (position 58 in the coding sequence) was based on a previous proteomic survey of mature N-termini of mitochondrial proteins in yeast^15^. To specifically observe the mature N-terminus of ALAS, we performed Edman degradation on ALAS purified from yeast cell extracts. ALAS appeared as a single processed species by Western blot (with possible trace uncleaved preprotein remaining, Fig. S2A). The N-terminal sequence of this species corresponded to cleavage after residue 34 of the preprotein (Δ34-ALAS), indicating that an additional 23 amino acids are retained by the mature protein beyond the previously detected species (Δ57-ALAS) (Fig. 2A, Fig. S2B). One possible source of this discrepancy is that the slower isolation procedure used to globally identify mitochondrial N-termini may have resulted in additional processing to yield Δ57-ALAS.

**Figure 2.**
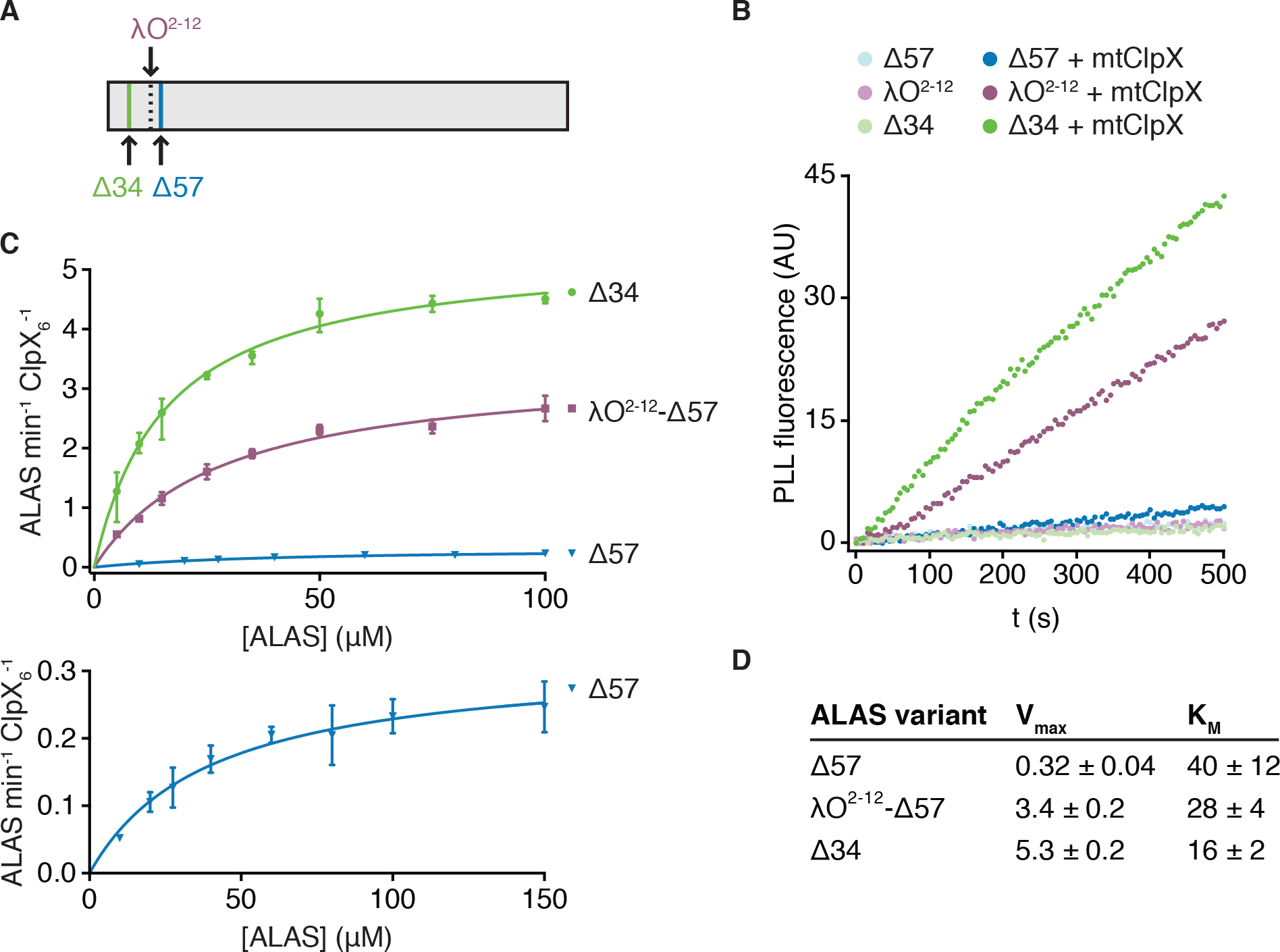
ClpX relies on an unstructured N-terminal extension for rapid activation of ALAS. (**A**) Edman degradation of ALAS identified a single N-terminus corresponding to amino acid 35 of the preprotein. A c-terminally His_7_ tag was integrated at the genomic locus encoding ALAS and the tagged protein was purified from yeast cell extract by Ni-NTA affinity. **(B)** Fluorescence traces representing PLP binding to 5 μM ALAS variants ± 2 μM mtClpX hexamer, monitored as in Fig. 1D. **(C)** The rates at which mtClpX stimulated PLP binding to Δ57-ALAS (blue, n = 2), λO^2-12^-Δ57-ALAS (purple, n = 2), or Δ34-ALAS (green, n = 3) are plotted as a function of ALAS monomer concentration. The lower graph shows the same fitted data for Δ57-ALAS as displayed in the upper graph with a smaller y-axis scale and an additional concentration (150 μM) not monitored for the other variants. mtClpX, when included, was present at 0.5 μM hexamer (λO^2-12^- and Δ34-ALAS variants) or 1 μM hexamer (Δ57-ALAS). PLP was included at 150 μM. Rates were extracted by linear fits to the early phase of PLP-binding fluorescence traces; the rate of mtClpX action was determined by subtracting the ALAS-alone PLP binding rate from the mtClpX-stimulated rate. Curves represent fits of the Michaelis-Menten equation (Y = V_max_*X/(K_m_ + X) to the data using Prism. **(D)** Kinetic parameters for mtClpX action on ALAS variants, extracted from fits in **(C)**. Standard error of the fit is stated.

We tested the effect of this natural N-terminal extension of ALAS on mtClpX activation of ALAS and found that mtClpX activated Δ34-ALAS even more rapidly than λO^2-12^-ALAS (Fig. 2B). Extension of the N-terminus of ALAS, either with its natural sequence (34-57) or with λO^2-12^, primarily stimulated the maximal rate of activation (a nearly twenty-fold increase in V_max_ by extension with the natural sequence) with a small effect on the K_M_ (Fig. 2B-D). The sequences of the λO^2-12^ tag and of the N-terminal extension of ALAS (amino acids 35-57) are not similar (Fig. S2B), suggesting that the role of this extension may be sequence independent. We did not detect any additional binding sequences in a peptide array including the entire preprotein sequence of ALAS, additionally supporting a sequence-independent interaction (Fig. S1D). This extended sequence has a dynamic or disordered structure; the N-terminus of Δ34-ALAS to the beginning of α1 (residue 71) exhibited near-instantaneous completion of deuterium uptake in hydrogen-deuterium exchange experiments (Fig. S2C; discussed further below). Therefore, mtClpX is a more potent activator of ALAS than we had previously appreciated, facilitated by a flexible element at the natural mature N-terminus of ALAS.

### Multiple sequence-specific contacts direct mtClpX action on ALAS

To identify residues that recruit mtClpX and position it on ALAS for unfolding and activation, we scanned the peptide sequences that bound mtClpX (Fig. 1A) with alanine and aspartate on an additional peptide array probed with mtClpX (Fig S3). In this array, we identified point mutations in each peptide that caused near-complete loss of mtClpX binding, with the exception of the most C-terminal peptide (Fig. S3). Most residues sensitive to mutation were perturbed by both alanine and aspartate. Several bulky hydrophobic amino acids (leucine, phenylalanine, and tyrosine) were highly represented in positions important for mtClpX interaction. This side-chain preference differed from the characterized preference of *E. coli* ClpX, for which basic residues are often important in substrate recognition^16^. The mtClpX-preferred residues we observe are also notable in that they comprise the set of signal residues described in the mitochondrial N-end rule^15^.

To test the role of these binding sequences in ALAS binding and activation, we mutated several residues that were important for mtClpX binding to ALAS-derived peptides in Δ34-ALAS. Alanine substitution at positions in α1 (F71A, Y73A, and the double substitution F71A/Y73A) reduced coprecipitation with mtClpX (Fig. 3A,B), and increased the K_M_ for activation by mtClpX (Fig. 3C-E), consistent with α1 functioning as a sequence-specific tag that recruits mtClpX to begin pulling on ALAS.

**Figure 3.**
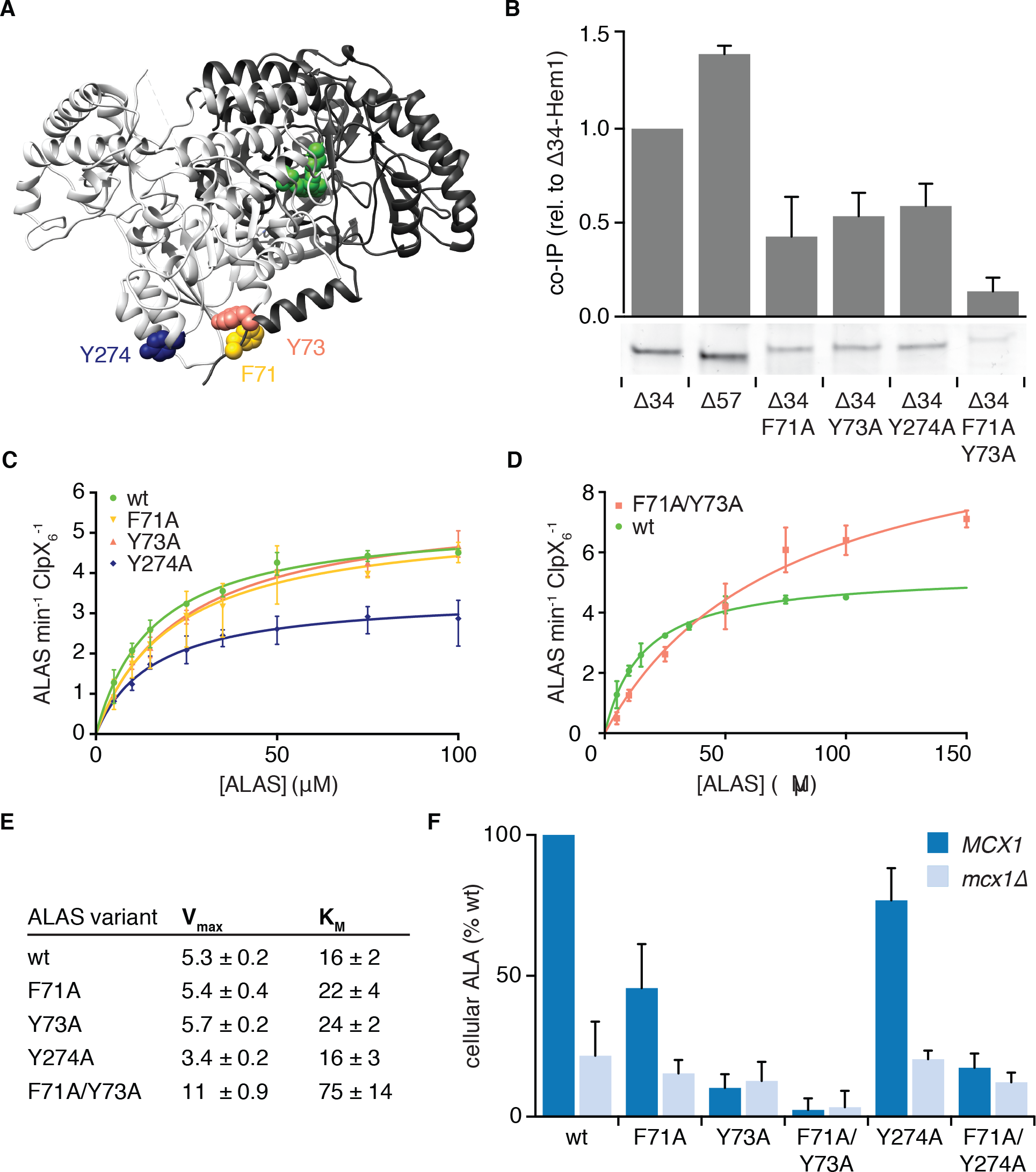
Multiple sequence-specific contacts direct mtClpX action on ALAS. (**A**). Position of mutations that perturb mtClpX binding, mapped on one face of the structure of ALAS (PDB: 5TXR) (F71 in yellow, Y73 in orange, and Y274 in dark blue). PLP is depicted in green. **(B)** Coimmunoprecipitation of ALAS variants with mtClpX^E206Q^-3xFLAG. 1 μM ALAS (monomer) and 0.5 μM mtClpX^EQ^-3xFLAG (hexamer) were incubated on ice with anti-FLAG antibody-conjuaged magnetic beads. Coprecipitating proteins were eluted with 3xFLAG peptide. Eluted proteins were separated by SDS-PAGE and stained with Sypro Red. The bar graph above each lane of the gel represents the average intensity of the band from three independent experiements, normalized to wt Δ34-ALAS; error bars represent SD. **(C-D)** The rates at which mtClpX stimulated PLP binding to indicated ALAS variants are plotted as a function of ALAS monomer concentration; rates were determined and fit to the Michaelis-Menten equation as in Fig. 2B. mtClpX was present at 0.5 μM hexamer (C) or 1 μM hexamer (D). Wildtype ALAS (Δ34-ALAS) data represented in Fig. 2C is replotted in both panels. N = 3 for all variants. **(E)** Chart of parameters extracted from fits in (C-D) as in Fig 2C. Standard error of the fit is stated. **(F)** The levels of ALA in extracts from yeast strains harboring the indicated mutations in ALAS (*HEM1* gene), with + (*MCX1*) or - (*mcx1Δ*) the gene encoding mtClpX, were measured by colorimetric assay with modified Ehrlich’s reagent. P < 0.001 for reduced ALA production in *MCX1* strains by all mutations displayed in *HEM1*; p = 0.05 for reduced ALA production in *mcx1Δ hem1*^*F71A/Y73A*^, n ≥ 3 for all strains; error bars represent SD.

The double substitution variant (ALAS^F71A/Y73A^) additionally had an increased rate of spontaneous PLP binding (Fig. S4A) and had an increased V_max_ for mtClpX-stimulated PLP binding (Fig. 3D,E). These residues are positioned at the dimer interface: Y73 directly contacts the other protomer through hydrogen bonding, and F71A is positioned for a CH-pi interaction with P246 (Fig. S4B). Although F71A and Y73A single variants retained synthase activity similar to wildtype, the double variant ALAS^F71A/Y73A^ retained only ~10-20% of wildtype activity (Fig. S4C). Removing contacts between the protomers through these mutations likely increases the solvent accessibility of the active site, thus increasing the spontaneous PLP binding rate of ALAS at the expense of active site function. Because the rate of unfolding of protein substrates by ClpX homologs is proportional to the mechanical stability of their fold, this reduced contact may be responsible for the increased V_max_ of mtClpX action on ALAS^F71A/Y73A^. These residues in α1 thus participate both in mtClpX recognition and in stabilizing contacts across the dimer interface that prevent spontaneous PLP exchange and promote activity.

In contrast to mutations in the N-terminal helix, a mutation at the surface of the mtClpX-interacting sequence cluster across the dimer interface from this helix, Y274A, did not perturb the K_M_ for mtClpX activation of ALAS (although it did reduce coprecipitation (Fig. 3B)). The Y274A mutation instead reduced the V_max_ of mtClpX activation (Fig. 3D, E). Y274 and other residues in this peptide cluster therefore may primarily interact with mtClpX to promote its rapid processing of ALAS.

To test if the contacts between ALAS and mtClpX we had identified were important for mtClpX to support ALA production in vivo, we introduced alanine mutations of the contact residues into the single genomic copy of ALAS in yeast (*HEM1)* in a wildtype or mtClpX-null (*mcx1Δ*) background. Within α1, F71A reduced and Y73A abolished mtClpX stimulation of ALA production in vivo (Fig. 3F). Combined mutation of both positions (*hem1*^*F71A/Y73A*^) reduced ALA production further, in agreement with the low activity of the corresponding protein variant in vitro. This protein variant was also reduced in abundance (Fig. S4D). Deletion of mtClpX in this double mutant caused no further reduction in ALA production. Mutation of the trans-protomer contact for mtClpX, Y274, modestly reduced mtClpX-dependent ALA production in vivo. Combined mutation of this contact and of one mtClpX α1 contact, F71 (*hem1*^*F71A/Y274A*^) abolished mtClpX stimulation of ALA production. The concordance between the effect of mutations in ALAS on its activation by mtClpX in vitro and mtClpX-dependent ALA production in vivo support the compound mtClpX-binding site spanned by these mutations as the physiological site at which mtClpX recognizes and initiates unfolding of ALAS to promote heme biosynthesis.

### mtClpX unfolds a defined region of ALAS that gates the active site

How does mtClpX unfolding from the N-terminus activate ALAS? We used hydrogen-deuterium exchange (HX) coupled with mass spectrometry (MS) to monitor conformational changes in Δ34-ALAS undergoing remodeling by mtClpX. ALAS conformation overall was very stable in the absence of mtClpX. Most of ALAS exhibited either no measurable deuteration or gradual deuteration over the 60-minute observation period, characteristic of a tightly folded protein (Fig. S2C and Supplemental File 1). The peptides in the N-terminal region leading up to the beginning of α1, however, became deuterated to a high level within seconds, consistent with this region behaving as a flexible, disordered element that mtClpX could use as a grip site.

In the presence of mtClpX, deuteration was increased in a defined region of ALAS over time (Fig. 4A, B; Supplemental File 1), consistent with mtClpX enacting a limited, specific conformational change rather than globally unfolding ALAS. mtClpX-induced deuterium uptake was localized to α1, the β sheet that connects α1 to the active site (β1-3) and the following active-site loop and α2, and several non-contiguous peptides surrounding the active site of ALAS (Fig. 4A, B). The mass spectra of most of these peptides exhibited clear EX1 kinetics^17^ (Fig. 4A, right panel, indicated with the half-life of the EX1 unfolding event, t_1/2_), consistent with cooperative unfolding within the peptide. This increased exposure was ATP-dependent and thus due to active unfolding by mtClpX (Fig. S5B and Supplemental File 1).

**Figure 4.**
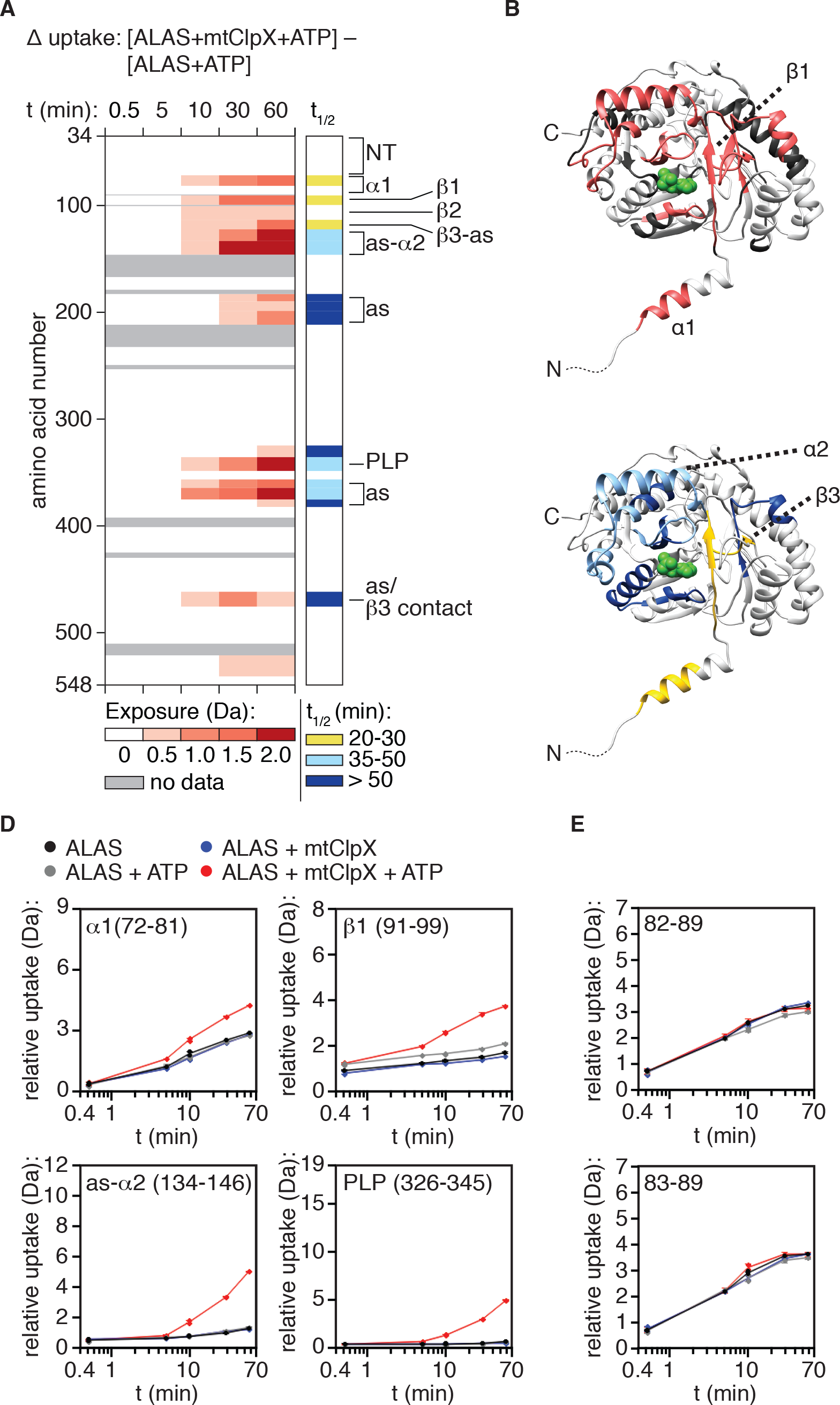
ClpX remodels a limited region of ALAS that extends from its N-terminal binding site to the active site of ALAS. **(A)** Left panel: mtClpX-induced deuterium uptake in ALAS (difference in deuterium uptake of ALAS with mtClpX and ATP present and ALAS with ATP only). A set of linear, nonoverlapping peptides is shown. Legend indicates minimum value of induced deuterium uptake for each color. See also Supplemental File 1 for difference maps of all peptides monitored. Peptides used in the linear map are indicated in this file. Right panel: half-life (t_1/2_) of mtClpX-induced exchange for peptides in which EX1 kinetics could be clearly assigned. To the right of both panels, the correspondence of structural and functional elements of interest in ALAS with detected peptides are indicated as follows: NT: (flexible N-terminus), 35-52, 53-71; α1: 72-81, 82-89; β1: 91-99; β2, 101-113; β3-A (β3 + active site-proximal sequence): 114-122; as-α2, 125-133, 134-152; as: 184-190, 191-199, 201-212; PLP (PLP-binding active site lysine): 326-345; as: 357-363, 364-374; as/β3 contact (tertiary structure contact with β3): 461-476. **(B)** mtClpX-induced deuterium uptake above 0.5 Da at 10 min, is mapped in salmon on the structure of ALAS (PDB: 5TXR). One protomer of the dimer is displayed; PLP is depicted in green. **(C)** t_1/2_ of the mtClpX-induced EX1 deuterium signatures from (A) mapped on one protomer of the ALAS dimer as in (B). Colors correspond to t_1/2_ as in the right panel of(A). **(D)** Plots of deuterium uptake over time ± ATP, ± ClpX for selected peptides (amino acid coordinates in parentheses) from ALAS that exhibit mtClpX-induced deuterium uptake **(E)** Deuterium uptake plots of peptides (amino acid coordinates indicated) in the gap in mtClpX-induced exposure of ALAS at the α1-β1 junction. Colors in plots correspond to conditions as indicated in legend in (D).

The β-sheet that mtClpX exposes partially shields the active site and contacts other active-site-proximal elements, suggesting that it could gate PLP access. To test whether exchange in this region was the direct result of mtClpX remodeling or due to PLP or PLP-oxime loss, we tested the effect of mtClpX on deuterium uptake of both PLP-depleted and holoenzyme preparations of ALAS, and observed a nearly identical uptake profile (Fig. S5C and Supplemental File 1). Inclusion of PLP in the exchange reaction only slightly suppressed some mtClpX-induced exposure immediately adjacent to bound PLP (Fig. S5D,E; see Supplemental File 1 for data from longer peptides with clear PLP-dependent protection), indicating that the increased exposure observed was due in most part to remodeling by mtClpX.

Two non-homogeneous features in the linear path of mtClpX-induced exchange suggest that mtClpX may not unfold along the entire length of this path. First, the unfolding half-life (t_1/2_)^18^ for mtClpX-induced exchange in these elements is not uniform: exchange in α1 and β1-3 has a shorter t_1/2_ than the exchange further along this path (as-α2 indicated in Fig. 1A, C) and in the non-contiguous peptides surrounding the active site (Fig. 4A right panel; Fig. 4C). These regions therefore may be exposed as a secondary and/or lower-probability consequence of mtClpX remodeling of α1 and β1-3, giving rise to a longer t_1/2_ for deuterium uptake. Second, a short sequence at the junction between α1 and β1 (82-89) exhibited no mtClpX-induced exchange (Fig. 4D,E). A possible explanation for this observation is that mtClpX arrests when its pore reaches this site. Occupancy by mtClpX could confer similar protection from solvent as the native structure of ALAS while regions further into the protein are deprotected by conformational change propagating from this site.

Together with our mapping of mtClpX binding sites, these data indicate a path of unfolding from the N-terminus of ALAS through some portion of the first β sheet. Unfolding along this path could thus gate access of PLP to the active site.

### Opening the β1-3 gate is required for ALAS activation

To test if mtClpX unfolding of ALAS from the N-terminus through α1 and some or all of the following β sheet is required to stimulate cofactor binding, we designed cysteine pairs flanking this path, such that formation of a disulfide bond between these pairs would block the linear path for ClpX unfolding (Fig. 5A). No disulfide bonds between the native cysteines in ALAS were predicted; in agreement with this prediction, nearly all wildtype ALAS protein remained as a monomer after oxidation (Fig. 5B). mtClpX stimulated oxidized and reduced ALAS to bind PLP at a similar rate (Fig. 5C, D), providing a clean background against which to observe perturbation by introduced disulfide pairs. We designed one cysteine pair (positions 68 and 243) to link the N-terminus of α1 to an adjacent site in the other protomer and a second pair (positions 88 and 427) to link the α1-β2 junction with an adjacent site in the same protomer (Fig. 5A). ALAS^68×243^ and ALAS^88×427^ both ran as monomers when reduced, but upon oxidation exhibited a near-complete shift in mobility. Oxidized ALAS^68×243^ shifted to an apparent higher molecular weight, approximately twice that of ALAS monomer, consistent with the cross-protomer disulfide bond. Oxidized ALAS^88×427^ shifted to a slightly faster-migrating species, consistent with the intramolecular crosslink between these residues (Fig. 5B).

**Figure 5.**
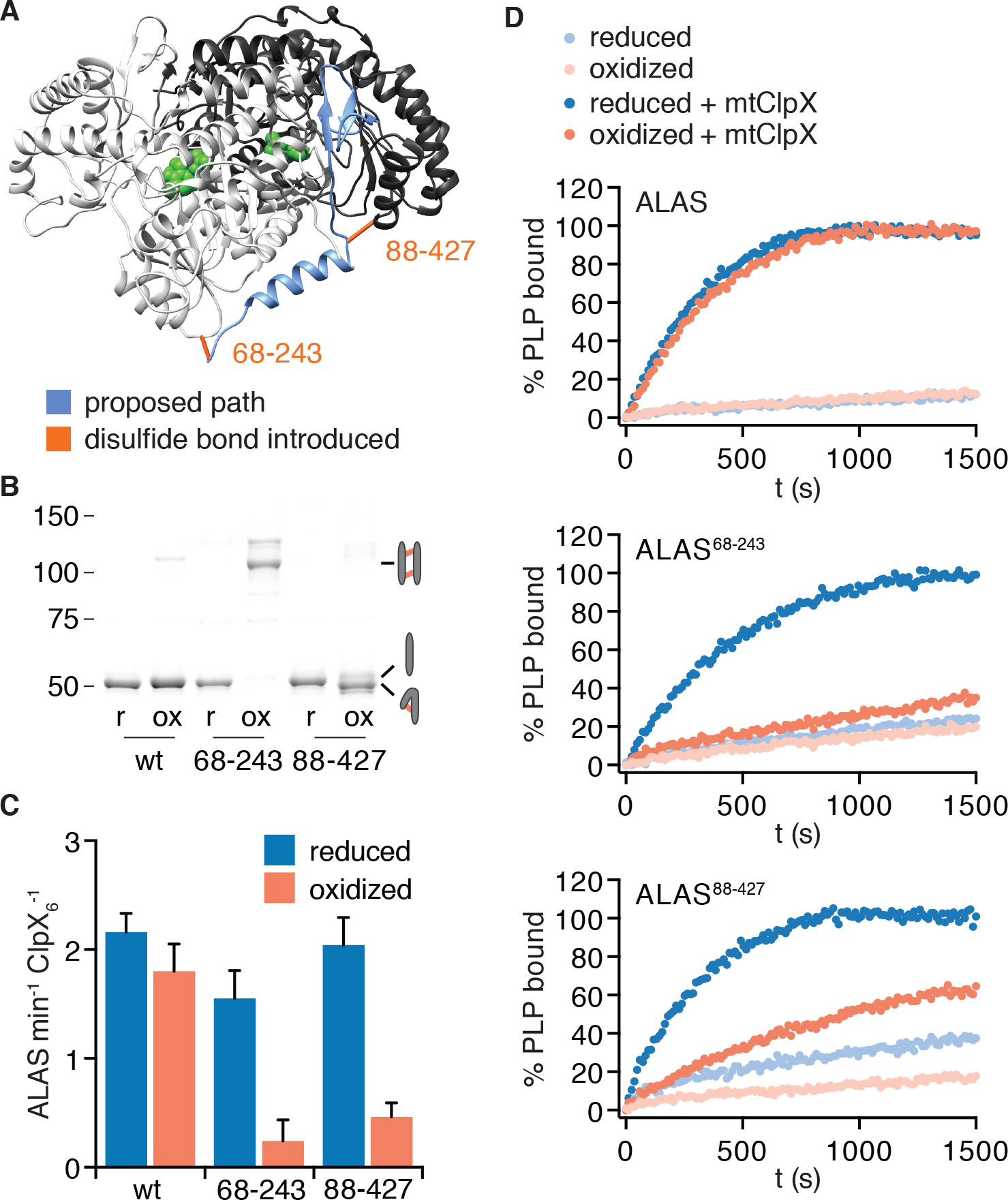
Unfolding of N-terminal secondary structure is required for mtClpX to activate ALAS. **(A)** The proposed path of mtClpX unfolding of ALAS to the active site is indicated in blue on The positions of cysteine pairs (indicated by amino acid number) introduced into ALAS are indicated by orange bonds on the structure of yeast ALAS (PDB: 5TXR). **(B)** Samples of ALAS cysteine variants (wt, 68-243, and 88-427) were separated by nonreducing SDS-PAGE and stained with Sypro Red, after oxidation (induced by addition of copper phenanthroline) and after reduction of oxidized samples with TCEP. Molecular weight marker (MW) sizes are indicated in kD. Cartoons indicate expected migration of species with intermolecular crosslinks (top right), uncrosslinked (middle right), and intramolecular crosslinks (bottom right). **(C)** mtClpX stimulation of PLP binding to indicated ALAS variants after oxidation and after re-reduction, monitored by fluorescence (ex. 434 nm, em. 515 nm). ALAS variants were present at 10 lM (monomer), PLP at 150 μM, ATP at 2 mM + regenerating system, and mtClpX, when included, at 0.5 XM (hexamer). Error bars represent SD from three independent experiments. p = 0.002 for suppression of mtClpX action by oxidation of ALAS^68×243^ and 10^-4^ **for** ALAS^88×427^ (compared to reduced form of same variant). **(D)** Representative traces of PLP binding to ALAS variants. 100% PLP-bound value was set to observed plateau of PLP fluorescence.

mtClpX action upon both oxidized variants was strongly attenuated compared to wildtype (Fig. 5C, D). Upon re-reduction of the crosslink, mtClpX induced both cysteine-pair variants of ALAS to bind PLP at a similar rate as for wildtype ALAS, indicating that loss of mtClpX action on these variants was specifically perturbed by disulfide bond formation (Fig. 5C, D). Therefore, mtClpX must unfold from its N-terminal engagement site on ALAS and extract at least the first β strand of ALAS in order to activate ALAS.

## DISCUSSION

We sought to define the mechanism by which mitochondrial ClpX deploys its unfoldase activity to activate ALAS, the initiating enzyme in heme biosynthesis. Here, we have defined sequence and structural elements that direct mitochondrial ClpX to bind ALAS and unfold a substructure of the protein, gating PLP cofactor to the active site. Our findings describe the mechanism of biogenesis and repair of an essential mitochondrial enzyme. This mechanism involves a diversion of mtClpX from the complete, global unfolding that ClpX homologs are best understood to execute. Our data indicate that mtClpX stalls in a region that lacks the few features previously described to stall unfoldases, suggesting that a greater variety of signals between unfoldase and substrate can direct alternative outcomes. This non-canonical action of a ClpX homolog also suggests that partial unfolding may be deployed to maintain and regulate a larger and more diverse set of cellular proteins than previously appreciated.

The elements comprising the multivalent ALAS recognition site affect different parameters of mtClpX action, suggesting a model for how they cooperate to recruit mtClpX and promote unfolding of ALAS. First, a composite site that includes α1 and a region immediately across the ALAS dimer interface recruits mtClpX using sequence-specific contacts. Mutation of residues in this site decreased the avidity of the ALAS-mtClpX interaction and insertion of an unfolding-resistant domain between this site and the body of ALAS blocked activation by mtClpX. We propose that these contacts recruit mtClpX and position it to pull efficiently on the ALAS structure. Once recruited, mtClpX initiates unfolding from the unstructured N-terminal tail of ALAS that precedes α1. Initiation of unfolding did not depend on the sequence of this tail and its deletion primarily increased the V_max_ of mtClpX activation rather than the avidity of interaction. This suggests that the N-terminal tail of ALAS provides a readily available grip for the pore loops of mtClpX to initiate unfolding. This composite signal, a sequence-specific recognition site and an sequence-independent unstructured element, is similar to a code that has been observed for protein unfolding by the proteasomal unfoldases: ubiquitin conjugation confers recognition of a protein substrate, but a nearby unstructured element is important for efficient unfolding^19^.

Several lines of evidence indicate that the way mtClpX engages this site can influence the outcome of the ALAS-mtClpX interaction. A dominant mutation in mtClpX was identified as the causative mutation in a human erythropoietic protoporphryia that results from overproduction of heme precursors. This mutation attenuated mtClpX function in vitro and slowed ALAS protein turnover in vivo^20^. A heme-binding motif in ALAS separately was observed to mediate mtClpX-dependent ALAS turnover under heme-replete conditions in vivo^21^. This motif falls within the flexible N-terminal region extension that we here identify as a grip site for mtClpX. The N-terminus of ALAS may thus serve as a bifunctional connection to mtClpX, directing activation when heme concentration is low, or degradation when high heme concentrations drive heme binding at this motif.

How does unfolding by mtClpX activate ALAS? Our HX MS observations of ALAS undergoing remodeling by mtClpX identified a linear region of ALAS that is remodeled. This region consists of α1, a small β-sheet that shields PLP from solvent, an active site loop, and α2. This linear path also sets an upper limit for how far mtClpX translocates into the structure of ALAS. Disulfide bonds introduced along this path blocked ALAS activation, indicating that mtClpX must unfold at least through the α1-β1 junction and extract β1 to efficiently activate ALAS. The gap in the linear path of mtClpX-induced exposure at the α1-β1 junction, as well as the disjunction in the halftimes of mtClpX-induced exchange through the first β sheet and after, suggest that this site may be the approximate stopping point for unfolding and translocation. mtClpX-induced exposure further into the active site may be a secondary consequence of unfolding in the β sheet. By extracting this small portion of the otherwise intact ALAS structure, mtClpX thus opens a gate to the active site for entry of PLP and/or release of damaged PLP species.

This limited unfolding within the compact structure of ALAS poses a further mechanistic question: how is mtClpX directed to stop, mid-unfolding, and release ALAS? A few previous examples of partial unfolding by a AAA+ unfoldase as part of proteolytic processing have been characterized, including the DNA polymerase clamp loader DnaX by ClpXP in *C. crescentus* and the transcription factors NFϰB and Ci by the 26S proteasome^22,23^. For these substrates, as well as some engineered model substrates, the truncation point is between independent domains; the unfoldase is limited by encountering a mechanically stable domain and an immediately preceding sequence that appears to stall the unfoldase. These sequences are often (but not always) low complexity in some way; how they interact with an unfoldase to promote stalling is not known. The partial unfolding of ALAS we observe does not obviously follow this code. The stopping point for mtClpX appears to be in the middle of a compact fold, rather than in a linker between domains. Although ALAS contains some alanine repeats in its natural N-terminus, replacement with the non-repetitive λO^2-12^ sequence still supported robust stimulation of cofactor binding by mtClpX.

Unlike ClpX, the proteasomal unfoldases, and other AAA+ unfoldase generalists that act on a wide variety of substrates, some AAA+ unfoldases act as dedicated modulators of one client protein. Several of these specialists, such as Rca, which de-inhibits Rubisco, or Trip13, which inactivates the mitotic checkpoint by remodeling Mad2, carry out a limited unfolding of their client that, like mtClpX with ALAS, does not appear to follow the stalling sequence/stable domain code^24,25^. Other substrate features and their interactions with the unfoldase may similarly direct both specialist AAA+ unfoldases and generalist AAA+ unfoldases with specialized substrates. The close apposition of the mtClpX binding site and its apparent stopping point suggests that the multivalent interactions that we have determined to recruit mtClpX to ALAS may themselves contribute to limiting its translocation of ALAS as well. Future probing of the arrest points of ALAS-mtClpX and these dedicated client-unfoldase pairs will illuminate signals that allow protein unfoldases to direct precise and varied outcomes for their substrates.

## MATERIALS AND METHODS

### Protein purification

*S. cerevisiae* ALAS (Hem1) and mtClpX (Mcx1) and related variants were expressed and purified as described previously^5^, with modifications of reducing agent (1 mM DTT) and protease inhibitors (0.5 mM PMSF). PLP-depleted ALAS was prepared by incubation with 5 mM hydroxlamine in 25 mM HEPES pH7.6, 100 mM KCl, 10% glycerol, and 1 mM DTT overnight on ice, followed by gel filtration (Superdex 200), concentration, and snap-freezing in liquid nitrogen. *E. coli* ClpX was purified as described previously^26^

### Strain and plasmid construction

The previously described *S. cerevisiae* ALAS expression plasmid (pET28b-H_6_-SUMO-57-ALAS)^5^ was modified by round-the-horn PCR to extend the N-terminus (Δ34-ALAS, λO^2-12^-ALAS) and by quickchange PCR to make point mutations. For DHFR fusions of ALAS, PCR products containing the *M. musculus* DHFR coding sequence, ALAS, additional N-terminal sequences as indicated, and short flanking regions from pET28b-H_6_-SUMO-Δ57-ALAS were assembled by gap repair in pRS315 in *S. cerevisiae*, followed by subcloning of the assembled sequence into pET28b-H_6_-SUMO.

### ALA measurement

Overnight cultures of yeast grown at 30°C in synthetic complete medium (CSM + YNB, Sunrise Science Products) + 2% glucose were used to inoculate cultures to OD_600_ 0.1-0.15 in the same media formulation. All cultures were grown to OD_600_ 1.0-1.25 at 30°C with shaking at 220 rpm. The equivalent of 10 mL at OD_600_ 1.0 was harvested by filtration. Cell extracts were prepared and ALA was measured by colorimetric assay using modified Ehrlich’s reagent as previously described^5^, except yeast extracts were cleared by centrifugation at 21,000 *g* rather than 2000 *g*.

### Peptide arrays

SPOT arrays of 15-amino-acid peptides, C-terminally linked to a cellulose membrane, were synthesized by standard Fmoc techniques using a ResPep SL peptide synthesizer (Intavis). Arrays were incubated with gentle agitation in methanol (5 min) followed by TBS (3 × 5 min) and then blocking solution (TBST + 5% nonfat dry milk) (2 h). Blocked arrays were then incubated with 0.5 μM mtClpX^E206Q^-3xFLAG hexamer in 25 mM HEPES pH 7.6, 100 mM KCl, 5 mM MgCl_2_, 10% glycerol, 0.05% Triton X-100, and 5% nonfat dry milk (1 h), washed in blocking solution (3 × 5 min), incubated with 1 μg/mL mouse anti-FLAG M2 antibody (Sigma) in blocking solution (1 h), washed in blocking solution (3 × 5 min), incubated with goat anti-mouse IgG-alkaline phosphatase conjugate (Bio-Rad #1706520) diluted 1:3000 in blocking solution (30 min), washed 5 min in block, then washed in TBST (3 × 5 min). Protein binding to the array was then imaged using ECF reagent (GE Healthcare Life Sciences) with a Typhoon FLA9500 scanner (GE Healthcare Life Sciences).

### Edman sequencing/protein isolation

*S. cerevisiae* ALAS was isolated from yeast cell extract by means of a C-terminal His_7_ tag (strain JKY220). 1 L YPD was inoculated to OD_600_ 0.05 from a saturated overnight culture and grown to OD_600_ 1.0 (30°C, 220 rpm shaking). Cells were harvested by centrifugation (3500 *g*, 5 min) and washed once by centrifugation in lysis buffer (50 mM Tris, 500 mM NaCl, 10 mM imidazole, 10% glycerol). The cell pellet was snap-frozen, thawed in 15 mL lysis buffer supplemented at 1:200 with a protease inhibitor cocktail (Calbiochem EDTA-free protease inhibitor cocktail III), lysed by French press (25 kPa), supplemented with 0.5 mM PMSF, and cleared by centrifugation (30,000 *g*, 20 min, 4°C). The cleared lysate was incubated with 0.5 mL Ni-NTA agarose beads (Qiagen) for 1 h with rotation at 4°C. The lysate-bead slurry was drained over a column and washed with 50 mL lysis buffer supplemented with 20 mM imidazole and 0.5 mM PMSF. Bound protein was eluted in lysis buffer supplemented with 250 mM imidazole, precipitated with 10% TCA, resuspended in Laemmli buffer, separated by SDS-PAGE, and transferred to Immobilon^PSQ^ PVDF. After staining with Ponceau S to detect bound protein, a band corresponding to the tagged ALAS protein was excised and the N-terminal sequence was determined by Edman degradation at the Tufts University Core Facility (10 cycles, ABI 494 protein sequencer (Applied Biosystems)).

### Fluorimetry - PLP binding

PLP binding to ALAS was monitored by fluorescence (ex. 434 nm, em. 515 nm) in PD150 (25 mM HEPES pH 7.6, 150 mM KCl, 5 mM MgCl_2_, 10% glycerol), with ALAS, mtClpX and *E. coli* ClpX concentrations as described in individual experiments.

Reactions additionally contained 2 mM ATP, an ATP regenerating system (5 mM creatine phosphate and 50 μg/mL creatine kinase), and 150 μM PLP, with the exception of experiments performed with 5 μM ALAS, for which PLP was included at 50 μM. Fluorescence was monitored in a 384-well plate using a SpectraMax M5 microplate reader (Molecular Devices) or in a quartz cuvette using a Photon Technology International fluorimeter.

### ALAS activity assays

ALA synthase activity of purified ALAS variant holoenzymes was monitored with 3 μM ALAS in PD150 with 100 μM succinyl-CoA and 100 mM glycine in a total volume of 60 μL. ALA content was determined using a procedure adapted from ^27^ as follows. After incubation for 1 minute at 30°C, 3 volumes of a 6.7% trichloroacetic acid and 13.3 mM N-ethylmaleimide solution was added to precipitate protein and quench residual DTT, respectively. After 15 minute incubation on ice, solutions were centrifuged for 10 minutes at 21000 *g* at 4°C. 150 μL supernatant was mixed with 50 μL 8% acetylacetone in 2 M sodium acetate and heated at 90¼C for 10 min. After cooling for 5 minutes, 150 μL of the resulting solution was mixed with 150 μL of modified Ehrlich’s reagent (20 mg/mL 4-(dimethylamino)benzaldehyde in 84% glacial acetic acid, 14% perchloric acid (70% stock) in a 96-well clear polystyrene plate and incubated for 15 minutes at room temperature.

### Hydrogen-deuterium exchange and LC-MS analysis

9 μM ALAS monomer was incubated in a 15-fold excess of deuterated PD150 at 20 tC with 0.5 μM mtClpX and 2 mM ATP added where indicated. At prescribed labeling times, the labeling reaction was quenched with an equal volume of ice-cold 0.3 M potassium phosphate pH 2.1. 10 μL of the quenched mixture (containing 44 pmoles ALAS) were injected onto a Waters nanoAcquity with HDX technology for online pepsin digestion and UPLC peptide separation as previously described^28^, with the exception that a 5-35% gradient of acetonitrile over 12 min was used. A Waters Synapt G2-Si with ion mobility enabled was used for mass analysis. Peptides were identified with triplicate undeuterated samples of ALAS alone, and in complex with mtClpX and ATP as indicated, using Waters MS^E^ and Waters Protein Lynx Global Server (PLGS) 3.0. Peptide maps were generated and deuterium incorporation was analyzed using Waters DynamX 3.0 software. Comparison experiments were performed in duplicate, yielding the same result. One representative replicate is shown for each comparison. Only peptides found in both experiments are shown. No correction was made for back-exchange, as conditions between the compared samples were identical.

### Disulfide bond design and crosslinking

To engineer disulfide bonds in ALAS, cysteine positions were chosen using the Disulfide by Design 2.0 software (http://cptweb.cpt.wayne.edu/DbD2/index.php)^29^. To induce disulfide bond formation in ALAS cysteine-pair variants, proteins were buffer-exchanged (Zeba spin columns, Pierce) into PD150 and incubated with 20 μM CuSO_4_ and 60 μM 1,10-phenanthroline for 15 minutes at 22¼C. For analysis by SDS-PAGE, oxidized samples were quenched with 1 mM EDTA and 1 mM N-ethylmaleimide in nonreducing Laemmli buffer. For analysis of activity, samples were buffer-exchanged into PD150. To prepare reduced protein for analysis, oxidized and buffer-exchanged samples were incubated overnight with 50 mM TCEP, then exchanged into PD150.

### PLP-oxime measurements

To quantify PLP-oxime bound to ALAS, a 5 μM solution of PLP-depleted ALAS was re-supplemented with 5 mM hydroxylamine in 60 mM Na_2_HPO_4_ and 30 mM HCl was boiled for 5 min, cooled on ice and precipitated by addition of a 60% volume of 12% perchloric acid followed by centrifugation (21,000 x *g*, 5 min, 4¼C). The supernatant was neutralized by mixing with equal volume of 0.5 M Na_3_PO_4_ and fluorescence (excitation 380 nm; emission 460 nm) was monitored (adapted from a previous method for PLP quantitation^30^). Fluorescence values were converted to PLP-oxime concentration by comparison with hydroxylamine-treated PLP standards.

## ACKNOWLEDGEMENTS

We thank Thomas E. Wales for assistance with MS method development and thoughtful discussion, I. Levchencko for the synthesis of peptide arrays, and Michael Berne and the staff of the Tufts University Core Facility for performing Edman sequencing. This work was supported by National Institutes of Health grants DK115558 to T.A.B. and GM101135 to J.R.E and fellowship to J.R.K. (DK095726) and by the Howard Hughes Medical Institute. J.R.K. and T.A.B. are employees of the Howard Hughes Medical Instistute. J.R.E. acknowledges a research collaboration with the Waters Corporation.

---

**Figure S1.**
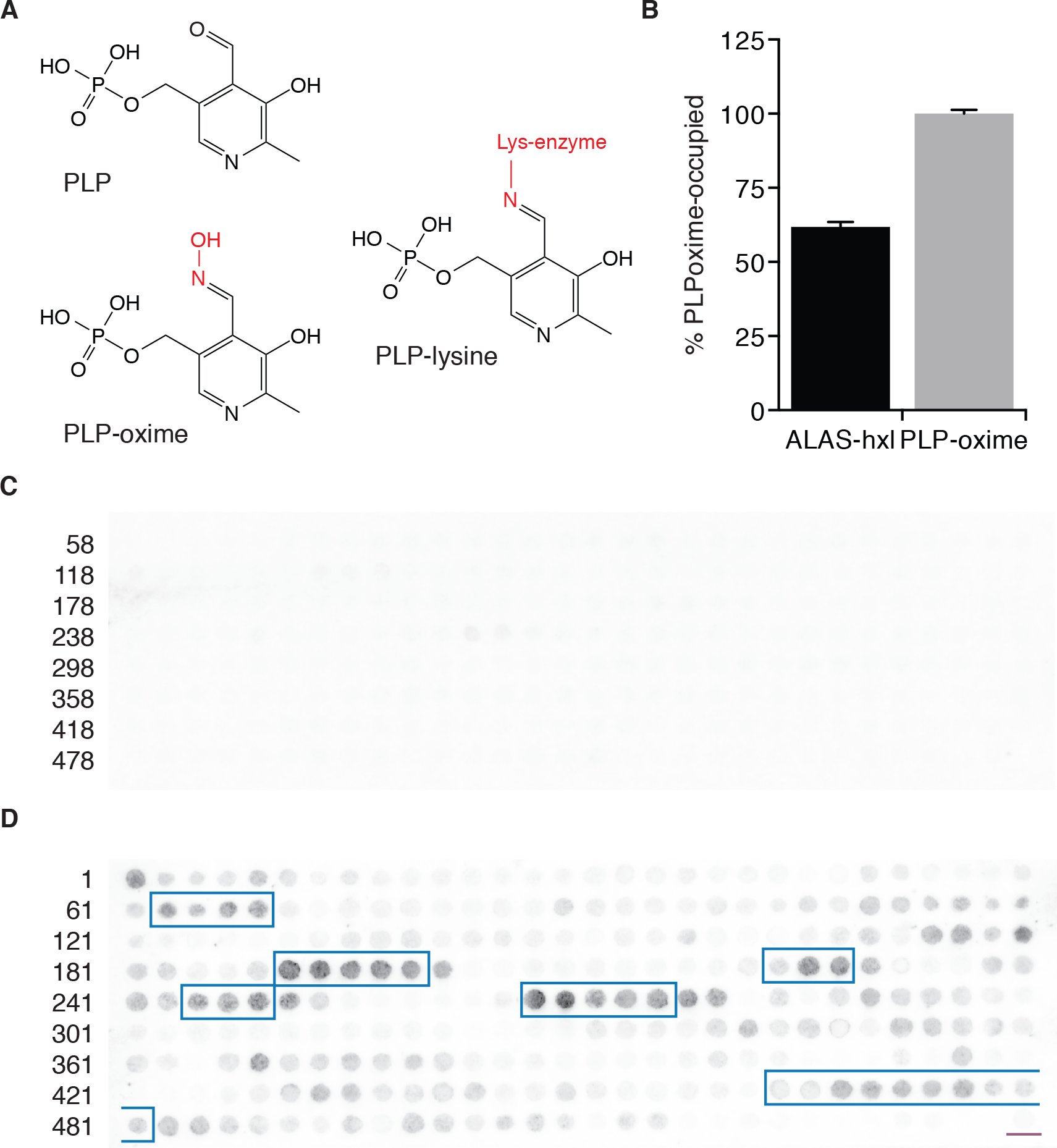
Related to Figure 1. **(A)** Pyridoxal phosphate and derivatives. Modifications of the pyridoxal aldehyde group caused by conjugation in an enzyme active site or by reaction with hydroxylamine are indicated in red. **(B)** PLP-oxime content of 10 μM hydoxylamine-treated ALAS, determined by fluorescence (ex. 380 nm, em. 460 nm) of the denatured and deproteinated solution. **(C)** Peptide array of Δ57-ALAS sequence (fifteen amino acid peptides linked C-terminally to a nylon membrane, shifted two amino acids towards the C-terminus with each spot, N-to C-terminus arrayed left-to-right, top-to-bottom), probed with anti-FLAG antibody followed by alkaline phosphatase-conjugated goat anti-mouse antibody and imaged by incubation with a fluorogenic alkaline phosphatase substrate (ECF, GE) on a Typhoon FLA9500 scanner (GE). Numbers indicate the first amino acid of the left-most peptide in each row. **(D) (E)** Peptide array of Δ34-ALAS sequence probed with mtClpX^E206Q^-3xFLAG and antibody as in (D). Numbers indicate the first amino acid of the left-most peptide in each row. λO^2-12^ peptide with alanine spacer (TNTAKILNFGRAAAA) is also included at the lower right (underlined in purple).

**Figure S2.**
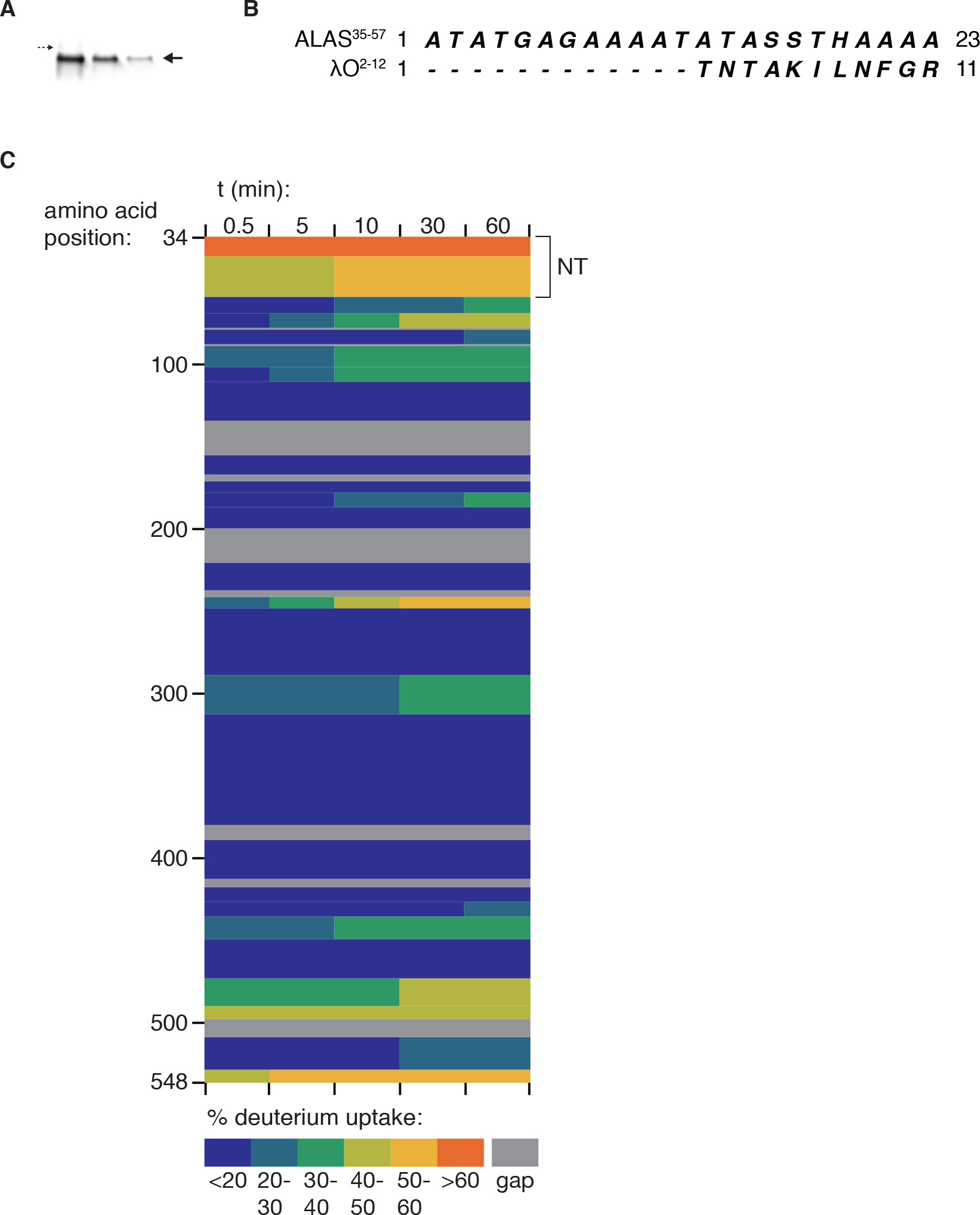
The extended, disordered mature N-terminus of ALAS, related to Figure 2. (**A**) Western blot (mouse anti-Myc) of ALAS-Myc-His_7_ purified from yeast cell extracts by Ni-NTA affinity. Lanes are samples from three sequential elutions. Large arrow at right indictates major mature species subjected to Edman degradation; small arrow at left represents indicates possible trace uncleaved preprotein. **(B)** Comparison of the sequence of the extension of the natural ALAS sequence beyond the previously annotated mature N-terminus (35-57) **(C)** Deuterium uptake in Δ34-ALAS, analyzed by HX MS. Only linear, non-overlapping peptides are shown; see Supplemental File 1 for deuterium uptake profiles of all peptides reproducibly detected.

**Figure S3.**
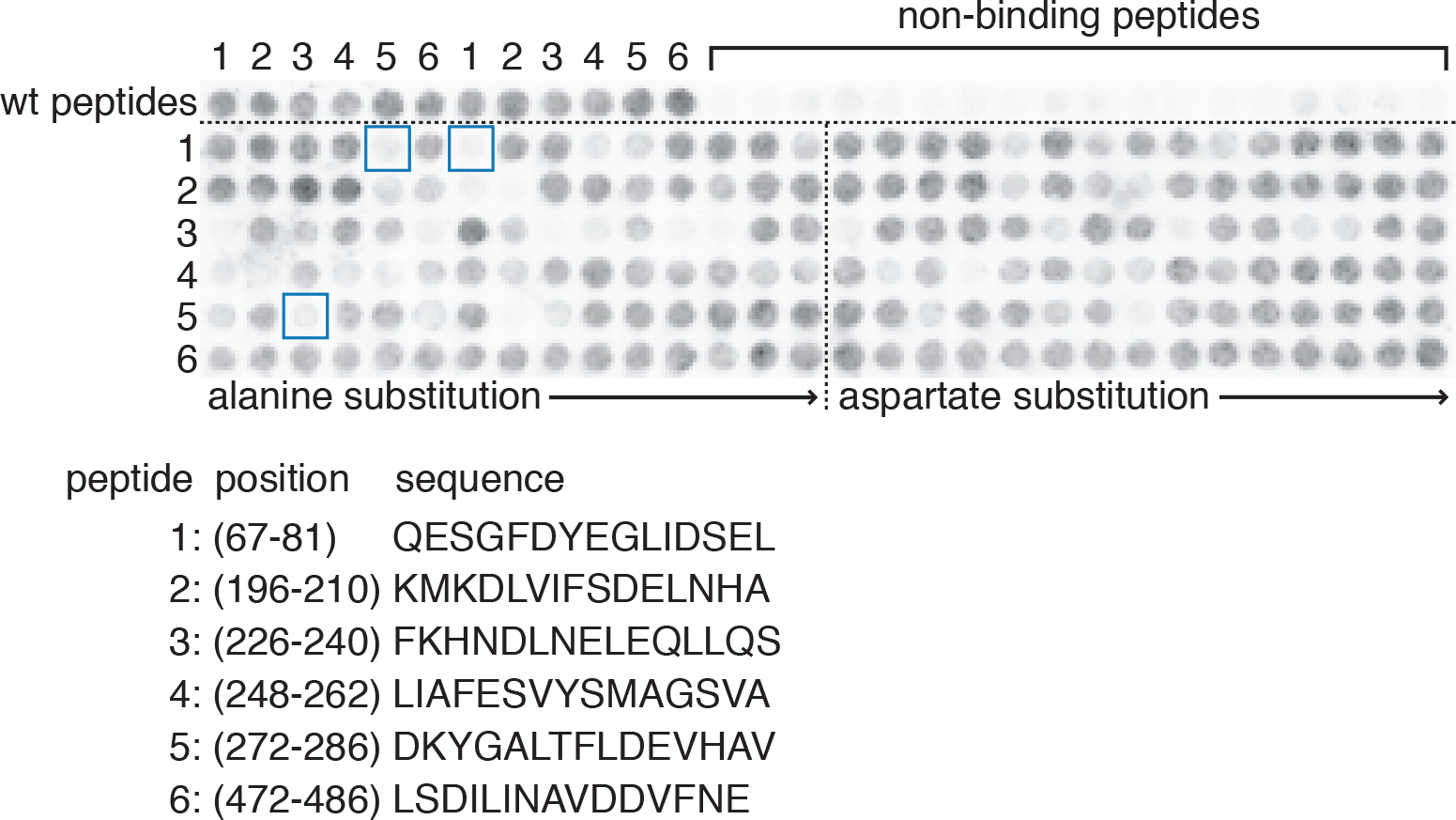
mtClpX-recognition sequences in ALAS, related to Figure 3. 15-amino-acid peptides derived from mtClpX-binding sequences determined in Fig. S2 were scanned at each position with alanine and aspartate in a peptide array. The top row of the array contains the wildtype peptides and non-binding peptides (peptides that mtClpX did not bind in Fig. S#2). Non-binding peptides are arrayed in the following series, repeated three times: LIDSELQKKRLDKSY (76-90), DSELQKKRLD-KSYRY (78-92), LEQLLQSYPKSVPKL (234-248), QLLQSYPKSVPKLIA (236-250), VRDPIVKQLEVSSGI (532-546), DPIVKQLEVSSGIKQ (534-548). Blue boxes represent ALAS protein variants analyzed.

**Figure S4.**
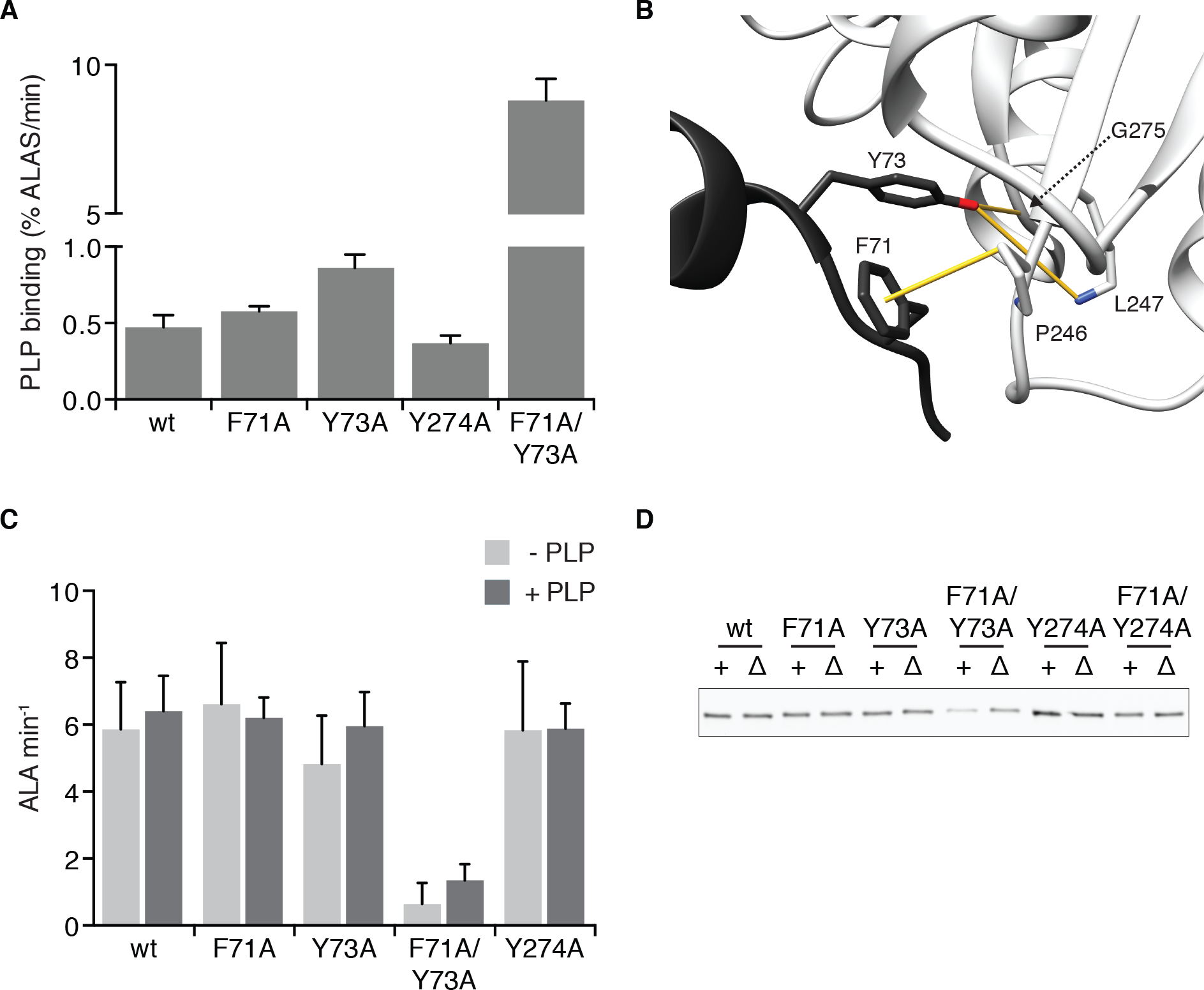
Structure and function of ClpX-interacting residues in ALAS, related to Figure 3. **(A)** Spontaneous binding rates of PLP to indicated ALAS variants. Rates were determined by linear fits to the pseudolinear early phase of PLP binding, measured by fluorescence (ex. 434 nm, em. 515 nm). ALAS variants were present at 10 μM, PLP at 150 μM, and ATP at 2 mM with a regenerating system. **(B)** CH-pi and H-bonding interactions (depicted as yellow bars) mediated by the side chains of ClpX-interacting residues F71 and Y73 (in protomer rendered in dark gray) with residues on the other protomer (rendered in light gray). **(C)** Activity of purified ALAS variants was monitored in vitro using modified Ehrlich’s reagent in a colorimetric assay (see Materials and Methods). Error bars represent SD of three independent assays; p < 0.01 for reduced ALAS^F71A/Y73A^ with or without PLP. **(D)** Relative protein levels of ALAS-3xMyc variants in vivo, determined by western blotting of yeast cell extracts with mouse anti-Myc antibody (9E10 clone) and IRDye^®^ 800CW Goat anti-rabbit IgG and visualized with an Odyssey scanner (Li-Cor).

**Figure S5.**
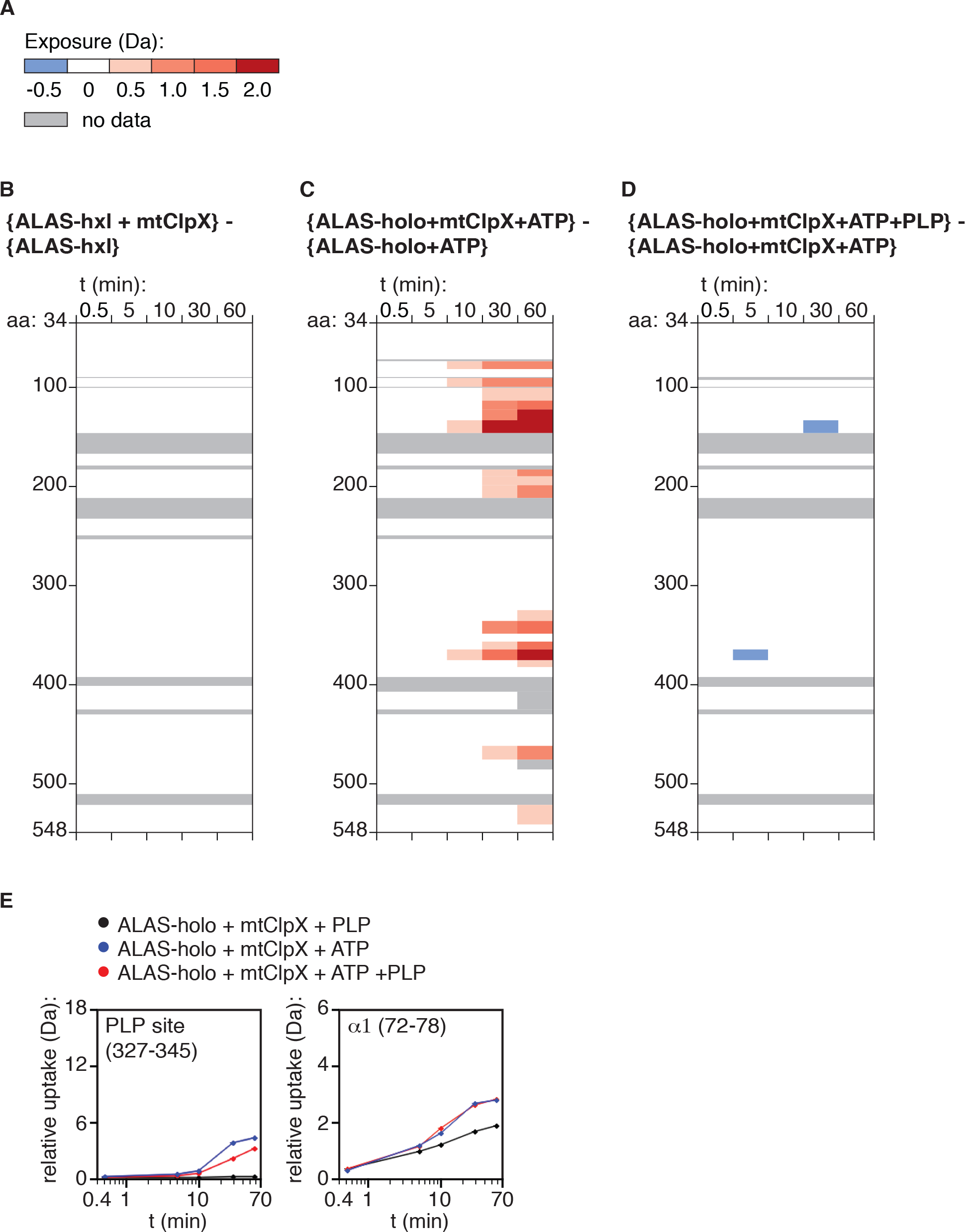
Deuterium uptake difference maps, related to Figure 4. **(A)** Legend for deuterium uptake difference maps in (B-D). Legend indicates minimum value of induced deuterium uptake for each color. **(B-D)** Difference maps comparing the states indicated composed of linear array of peptides corresponding to those shown in Fig. 4A. Deuterium uptake difference between the two states indicated was subtracted for each plot. See also Supplemental File 1 for difference maps of all peptides monitored. Specific peptides used in the linear maps are indicated in this file. **(E)** Plots of deuterium uptake over time for selected peptides that exhibit PLP-suppressed mtClpX-induced deuterium uptake (left) or PLP-insensitive mtClpX-induced deuterium uptake (right). 327-345 contains the PLP-bonding lysine (337); 72-78 lies in α1.

**Table S1.** *S. cerevisiae* strains used in this work. All strains were made in w303 mat a background (*MATa ade2-1 leu2-3 ura3 trp1-1 his3-11,15 can1-100 GAL psi+*)

| strain | genotype |
| --- | --- |
| JKY194 | *hem1Δ::6HA\*NAT* |
| JKY195 | *hem1Δ::6HA\*NAT mcx1Δ::KAN* |
| JKY196 | *hem1(F71A-Y73A)-3MYC\*TRP1* |
| JKY197 | *hem1(F71A-Y73A)-3MYC\*TRP1 mcx1Δ::KAN* |
| JKY198 | *HEM1-3MYC\*TRP1* |
| JKY199 | *HEM1-3MYC\*TRP1 mcx1Δ::KAN* |
| JKY220 | *hem1-MYC-7HIS\*KAN* |
| JKY229 | *hem1-F71A-3MYC\*TRP1* |
| JKY230 | *hem1-Y73A-3MYC\*TRP1* |
| JKY231 | *hem1-F71A-3MYC\*TRP1, mcx1Δ::KAN* |
| JKY232 | *hem1-Y73A-3MYC\*TRP1, mcx1Δ::KAN* |
| JKY236 | *hem1-Y274A-3MYC\*TRP1* |
| JKY237 | *hem1-Y274A-3MYC\*TRP1, mcx1Δ::KAN* |
| JKY238 | *hem1-F71A-Y274A-3MYC\*TRP1* |
| JKY239 | *hem1-F71A-Y274A-3MYC\*TRP1, mcx1Δ::KAN* |

**Supplemental File 1.** Deuterium uptake and difference values for all peptides monitored in ALAS.

